# NMDA receptor activation drives early synapse formation *in vivo*

**DOI:** 10.1101/2024.05.23.595343

**Authors:** Noah S. Leibold, Nathalie F. Higgs, Steffen Kandler, Adil Khan, Flavio Donato, Laura C. Andreae

## Abstract

The rules governing neural circuit formation in mammalian central nervous systems are poorly understood. NMDA receptors are involved in synaptic plasticity mechanisms in mature neurons, but their contribution to circuit formation and dendritic maturation remains controversial. Using pharmacological and genetic interventions to disrupt NMDA receptor signaling in hippocampal CA1 pyramidal neurons *in vitro* and *in vivo*, we identify an early critical window for a synapse-specific function in wiring Schaffer collateral connections and dendritic arborization. Through *in vivo* imaging, we show that NMDA receptors are frequently activated during early development and elicit minute-long calcium transients, which immediately precede the emergence of filopodia. These results demonstrate that NMDA receptors drive synapto- and dendritogenesis during development, challenging the view that these processes are primarily mediated by molecular cues.

## Introduction

Optimal development of synaptic connectivity is required for properly functional brain circuits, but the rules that drive precise wiring are not well understood. In the hippocampus, the development of appropriate circuit connectivity needs to allow efficient learning that will be essential for survival. Neuronal activity is crucial for the refinement and maintenance of circuits after they have formed, across multiple brain regions and cell types during development (*1-4*). However, its contribution to synapse formation *per se* and dendritic arborization remains controversial. Early work suggested a possible role for neurotransmission in synapse formation (*1, 5*), and there is evidence that NMDA receptors (NMDARs), which are expressed early in neuronal development (*6*) and are critical for synaptic plasticity in mature neurons (*7*), may facilitate dendritic branching (*8-10*). However, the discovery that the postnatal abolition of all excitatory transmission by genetic deletion of ionotropic glutamate receptors, including NMDARs, in hippocampal CA1 pyramidal neurons (PNs) was not associated with any changes to synapse densities or dendritic arbor structure cast doubt on this role (*11*). Indeed, recent evidence in zebrafish demonstrates that action potential firing is not required for the formation of circuits underlying visual behaviors (*12*). Hence, the current prevailing view is that synaptogenesis is driven by genetically encoded molecular cues, with activity playing a role only in subsequent refinement (*13*). Here we present evidence that reconciles these previous apparent contradictions. We demonstrate that NMDAR function is essential for synapse formation and dendritic elaboration in hippocampal CA1 PNs, but only in a prenatal critical developmental window, with no effects seen when blockade is initiated after birth. Furthermore, we find these effects are synaptic input specific. In addition, we confirm the lack of impact of neuronal firing, implicating the involvement of alternative sources for glutamate. Finally, by developing an approach to image developing CA1 PN dendrites *in vivo*, we discover exceptionally prolonged, localized dendritic calcium events that link to structural changes. Therefore, we find that NMDARs do contribute to circuit formation, but in a synapse-specific manner during early development.

### Prenatal window for NMDAR role in Schaffer collateral synapse formation *in vitro*

To identify whether timing might be important in the role of neuronal activity in synapse formation, we targeted two developmental time windows, before (embryonic day 18.5 (E18.5)) and after (postnatal day 1.5 (P1.5)) birth, using mouse hippocampal organotypic slices. Slices with neurons expressing GFP were treated with APV to inhibit NMDAR signaling or TTX to block action potential firing from day 1 to 3 *in vitro* (DIV) (Fig. 1A). We targeted the canonical Schaffer collateral (SC) projection from CA3 to CA1. Our imaging and analysis focused on the apical dendrites of CA1 PNs, with pre- and excitatory postsynaptic structures immunolabeled using bassoon and homer antibodies, respectively (Fig. 1, B and C). Synapses were identified through the colocalization of bassoon and homer puncta along GFP-labelled dendrites. Notably, TTX treatment showed no impact on synapse numbers at either stage, while blocking NMDARs with APV at E18.5 resulted in reduced homer and synapse densities (Fig. 1D). In contrast, postnatal blockade of NMDARs had no effect (Fig. 1E). These findings align with previous work indicating that action potential firing is not involved in synapse formation (*12*) but suggest that NMDARs play a role in the formation of SC synapses during an early, prenatal developmental time window *in vitro*.

**Fig. 1.**
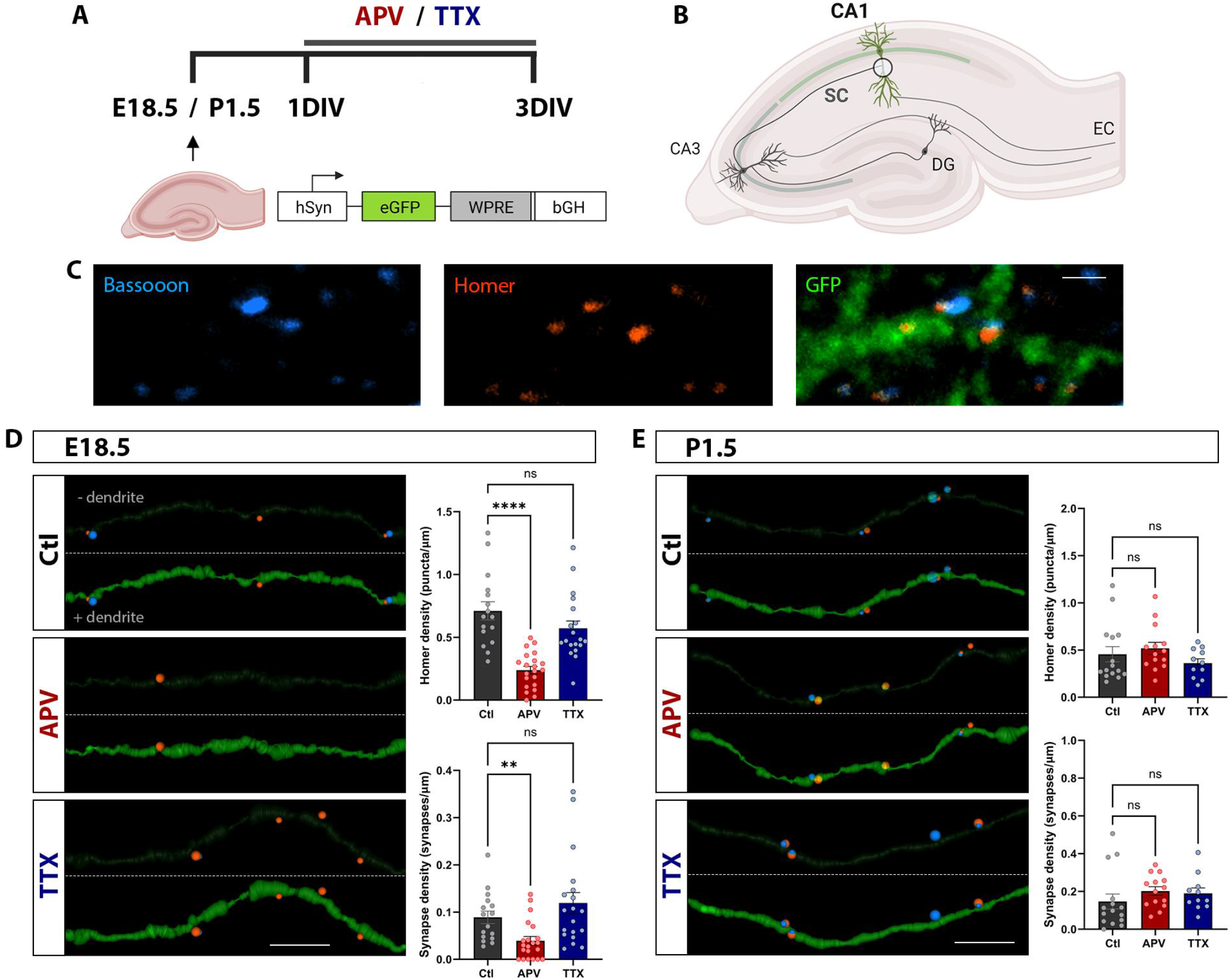
NMDARs contribute to synapse formation of CA1 SC synapses in hippocampal organotypic slices *in vitro*. (**A**) Schematic of experimental procedure: mouse hippocampal slices treated with AAV-GFP at E18.5 and P1.5 were cultured in APV or TTX between 1DIV and 3DIV. (**B**) Diagram of hippocampal anatomy: drug effects were assessed at CA1 PN apical SC synapses. (**C**) Immunostaining of bassoon and homer puncta with a GFP+ dendrite. Scale bar, 1 μm. (**D**) 3D reconstructions of dendrites and synapses in control (Ctl), APV- and TTX-treated neurons cultured at E18.5. Images are shown with (+ dendrite) and without dendrite (- dendrite) for better visualization of synapses. Comparison of homer densities (\*\*\*\**P* < 0.0001 for Ctl versus APV, *P* = 0.1914 for Ctl versus TTX; one-way ANOVA) and synapse densities (\*\**P* = 0.0097 for Ctl versus APV, *P* > 0.9999 for Ctl versus TTX; Kruskal-Wallis test), n = 16 Ctl, 21 APV, 19 TTX dendrites. Scale bar, 4 μm. (**E**) Reconstructions of dendrites and synapses in Ctl, APV- and TTX-treated neurons cultured at P1.5. Comparison of homer densities (*P* = 0.5359 for Ctl versus APV, *P* > 0.9999 for Ctl versus TTX; Kruskal-Wallis test) and synapse densities (*P* = 0.0863 for Ctl versus APV, *P* = 0.2650 for Ctl versus TTX; Kruskal-Wallis test). N = 15 Ctl, 14 APV, 11 TTX dendrites. Scale bar, 3 μm.

### NMDARs contribute to SC synapse formation *in vivo*

To assess the role of NMDARs in shaping hippocampal synaptic circuits and dendritic development *in vivo*, we selectively targeted hippocampal CA1 PNs for deletion of NMDAR function early in development by *in utero* injection of AAVs expressing Cre and a flexed reporter, tdTomato, at E14.5 in mice with a conditional deletion of the obligatory GluN1 subunit, *Grin1*^*fl/fl*^ (GluN1KO). Abolition of NMDAR function was tested by whole-cell patch clamp recordings of CA1 PNs at P3 – P4, targeted by a localized picospritzer puff of NMDAR agonists NMDA and glycine (Fig. S1, A and B). Holding cells in magnesium-free ACSF to unblock NMDARs, voltage clamp recordings of tdTomato+ neurons at -60 mV revealed an absence of NMDAR-mediated currents in GluN1KO cells (Fig. S1C). In contrast, *Grin1*^*fl/+*^ (Ctl) CA1 PNs exhibited robust NMDAR-mediated currents with comparable peak amplitudes to *Grin1*^*+/+*^ PNs (Fig. S1D). Similar findings persisted at P7 – P8, indicating a sustained loss of function during development (Fig. S2, A-C). In summary, we achieved a complete loss of NMDAR function by administering AAV-Cre early in embryonic development.

We then assessed the impact of this early loss of NMDAR function on synapses and morphology at P3. By titrating virus concentration we achieved sparse, yet bright labelling of CA1 PNs (Fig. 2, A and B). We labelled pre- and postsynapses by staining for bassoon and homer, respectively, and quantified their colocalization along reconstructed GFP+ dendrites to detect synapses (Fig. 2C). We examined SC projections from CA3 to basal and apical dendrites in CA1, and perforant path inputs from the entorhinal cortex (EC) to distal tuft dendrites (Fig. 2D). Intriguingly, GluN1KO neurons exhibited a reduction in excitatory synapse densities in basal and apical SC dendritic compartments, while in contrast, synapses from the EC remained unaffected by the loss of NMDAR function (Fig. 2E).

**Fig. 2.**
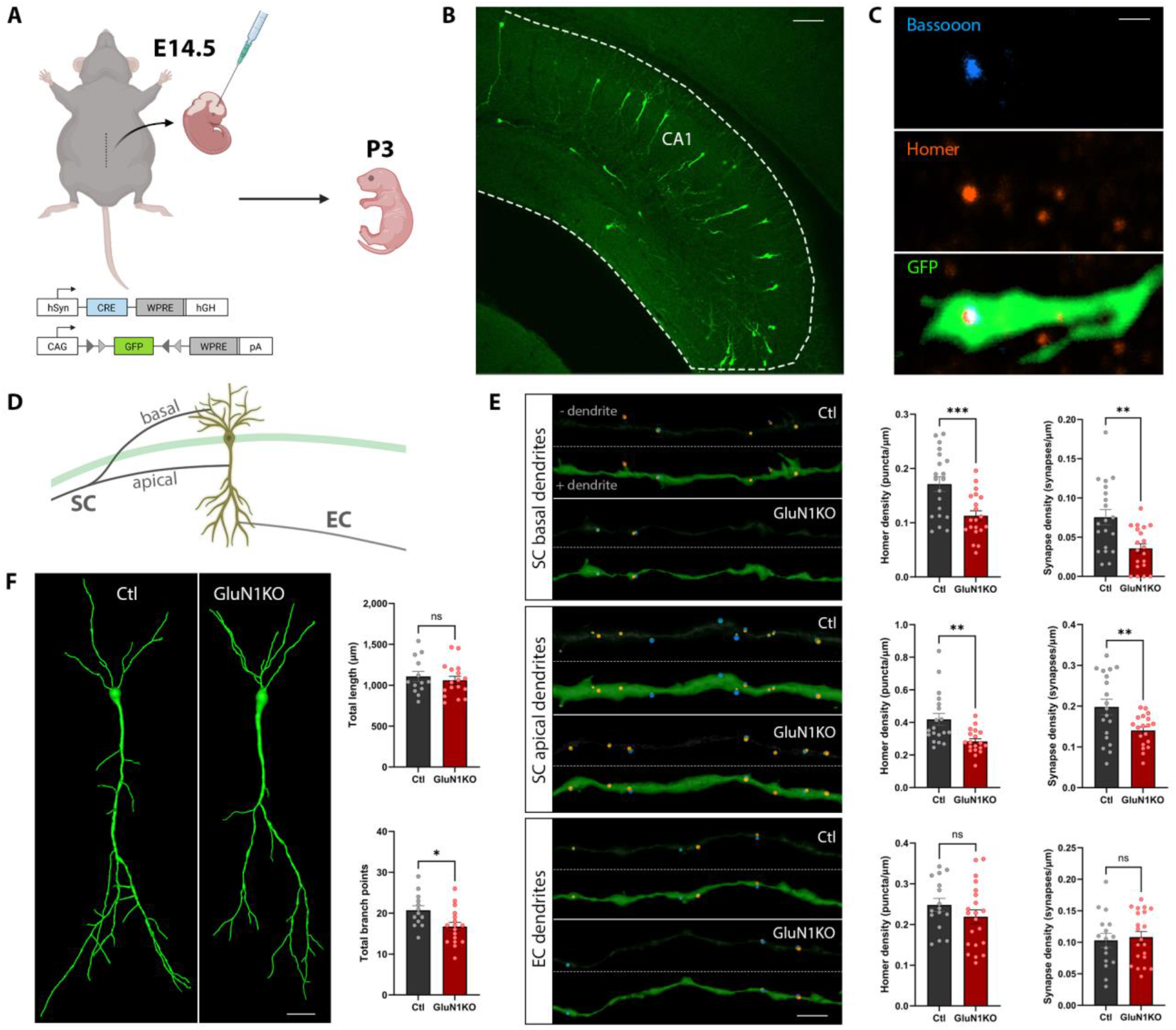
A synapse-specific role for NMDARs in guiding synapse formation in CA1 *in vivo*. (**A**) Schematic of E14.5 *in utero* viral injection of AAV-Cre and AAV-flex-GFP. (**B**) Confocal image of GFP+ CA1 PNs at P3 in a hippocampal slice. Scale bar, 100 μm. (**C**) Representative immunostaining images of bassoon and homer puncta adjacent to/within a GFP+ dendrite. Scale bar, 1 μm. (**D**) Diagram illustrating the analyzed synaptic connections received by CA1 PNs. (**E**) 3D reconstructions of dendrites and synapses. Top panel: synapses in CA1 basal dendrites (n = 20 Ctl cells from 5 mice; n = 20 GluN1KO cells from 5 mice). \*\*\**P* = 0.001 and \*\**P* = 0.0015 (unpaired *t* tests). Middle panel: synapses in CA1 proximal apical dendrites (n = 19 Ctl cells from 5 mice; n = 19 GluN1KO cells from 5 mice). Homer density: \*\**P* = 0.0013 (Mann-Whitney test). Synapse density: \*\**P* = 0.0098 (unpaired *t* test). Lower panel: synapses in CA1 apical tuft dendrites (n = 16 Ctl cells from 4 mice; n = 21 GluN1KO cells from 5 mice). Homer density: *P* = 0.2341 (unpaired *t* test). Synapse density: *P* = 0.7797 (Mann-Whitney test). Scale bar, 5 μm. (**F**) Whole cell 3D reconstructions of CA1 PNs. Quantification of total lengths and total number of branch points (n = 13 Ctl cells from 5 mice; n = 17 GluN1KO cells from 6 mice). *P* = 0.5543 and \**P* = 0.0167 (unpaired *t* tests). Scale bar, 5 μm.

While the data suggested a role for NMDARs in wiring synaptic circuits, there remained a possibility that their role could theoretically be to maintain pre-existing synapses. As a result, we aimed to delineate the course of synapse formation in Ctl CA1 PNs. Notably, synapses were almost absent in SC apical dendrites at P0 (Fig. S3A), whereas densities in EC dendrites showed no significant difference between P0 and P3 neurons (Fig. S3C). Furthermore, we observed a trend toward a reduction in SC synapses in GluN1KO cells at P0 (Fig. S3B), with no alteration in EC synapses (Fig. S3D).

To examine whether these differential effects between SC and EC synapses could be due to differences in NMDAR expression and function, we injected WT mice at E14.5 with an AAV expressing GCaMP6s and mRuby2, and focally applied NMDA and glycine to basal SC input sites in the stratum oriens (SO), apical SC synapses in the stratum radiatum (SR) and EC inputs in the stratum lacunosum moleculare (SLM) to acute hippocampal slices at P0 – P2 (Fig. S4A). Two-photon calcium imaging was used to quantify NMDAR responses. NMDARs were expressed and functional as early as P0 in SO, SR and SLM (Fig. S4, B-D) with no differences in GCaMP6s peak amplitudes between the compartments at any timepoint (Fig. S4E). These data provide evidence that NMDARs are expressed and functional during the time window of SC synapse formation. The absence of a synaptic phenotype in SLM at P3 cannot be explained by a lack of functional NMDARs in distal tuft dendrites, but may be related to the timing of their initial formation.

To assess impacts on dendritic morphology, we generated whole-cell reconstructions of CA1 PNs. While no difference was observed in total length, GluN1KO cells displayed fewer dendritic branch points (Fig. 2F). Collectively, our results support a role for NMDARs in wiring specific synaptic connections and promoting dendritic branching early in development.

### Dynamic dendritic branching patterns *in vivo*

Given that GluN1KO cells have fewer synapses and dendritic branch points at P3, we hypothesized that NMDARs may stabilize dendrites and promote branching and growth. We therefore tracked CA1 basal dendrites expressing a flexed reporter, tdTomato, at P3 – P4 *in vivo* (Fig. 3A). To assess the dynamic branching patterns of Ctl CA1 PNs early in development, we continuously imaged distal and proximal dendritic segments and tracked the displacement of branching filopodia (Fig. S5, A and B). Dendrites showed dynamic structural changes, with filopodia branching from distal dendrites exhibiting more pronounced alterations and covering a greater distance (Fig. S5, C and D). To compare structural changes between Ctl and GluN1KO cells, we imaged entire basal dendritic arbors of CA1 PNs at two timepoints, 0 and 2 hr, allowing for substantial structural changes to occur in the dendritic arbor. Basal arbors were reconstructed in 3D and aligned by dendrite in order to detect retracted and added dendrites (Fig. 3B, Fig. S6A). Compared with Ctl dendrites, which on average displayed an extension of branch length over this time period, GluN1KO dendrites showed significantly less extension, indeed tending to decrease in length (Fig. 3C). Moreover, GluN1KO cells showed an increase in retracted dendritic tips (Fig. 3D), and a tendency to add fewer branches (Fig. 3E). Considering that distal dendrites display more pronounced structural changes than proximal dendrites, we examined the location of retracted and added dendrites along the basal arbor. No differences were observed in spatial locations when comparing retracted with added dendrites in either Ctl or GluN1KO cells, along with total lengths (Fig. S6, B and C). Interestingly, Ctl cells had a higher proportion of added and retracted dendritic tips in distal dendritic regions compared to GluN1KO cells (Fig. 3, F and G). Additionally, a greater number of stable dendrites in Ctl neurons extended further into the neuropil environment when examining the pathlength from the soma to their distal tips (Fig. S6D). Collectively, these data support the role of NMDARs in stabilizing dendritic branches, thereby influencing dendritic complexity.

**Fig. 3.**
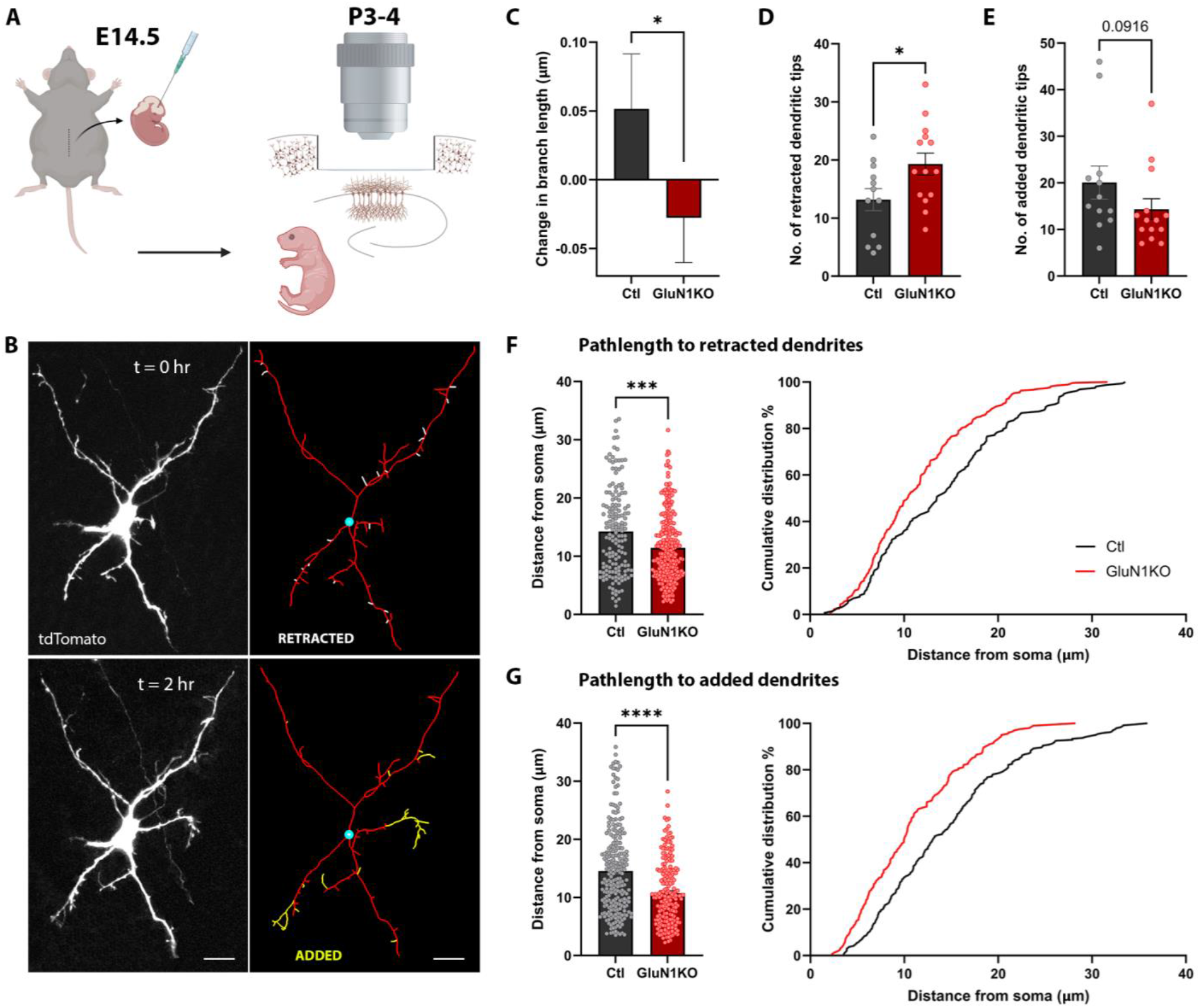
NMDARs stabilize and promote dendritic branching *in vivo*. (**A**) Schematic illustrating cranial window implant for optical access to CA1 PNs at P3-4. (**B**) Representative maximum image projections of an example tdTomato+ Ctl cell imaged at two timepoints, t = 0 hr and t = 2 hr, with reconstructions to show retracted (white) and added (yellow) branches. Scale bar, 5 μm. (**C**) Quantification of changes in branch length of all dendritic tips (n = 747 Ctl dendrites, 12 cells from 5 mice; n = 966 GluN1KO dendrites, 14 cells from 5 mice). \**P* = 0.0418 (Mann-Whitney test). (**D**) Total number of retracted dendritic tips per neuron (n = 12 Ctl cells from 5 mice; n = 14 GluN1KO cells from 5 mice). \**P* = 0.0322 (unpaired *t* test). (**E**) Total number of added dendritic tips per neuron (n = 12 Ctl cells from 5 mice; n = 14 GluN1KO cells from 5 mice). *P* = 0.0916 (Mann-Whitney test). (**F**) Measurement and cumulative distribution of pathlengths from the soma to the start point of retracted dendrites for all neurons (n = 158 Ctl lengths, 12 cells from 5 mice; n = 270 GluN1KO lengths, 14 cells from 5 mice). \*\*\**P* = 0.0002 (Mann-Whitney test). (**G**) Measurement and cumulative distribution of pathlengths from the soma to the start point of added dendrites for all neurons (n = 241 Ctl lengths, 12 cells from 5 mice; n = 201 GluN1KO lengths, 14 cells from 5 mice). \*\*\*\**P* < 0.0001 (Mann-Whitney test).

### Prolonged NMDAR-mediated dendritic calcium transients *in vivo*

Having observed structural alterations following loss of NMDAR function, we next assessed the functional impact at P3 – P4 *in vivo* by expressing the calcium reporter GCaMP8s in CA1 PNs. We were able to detect discrete, local dendritic calcium transients (Fig. 4, A and B). We observed a decrease in the mean frequency of local calcium transients in GluN1KO neurons compared to Ctl, with no difference in mean amplitude (Fig. 4, C-E, Fig. S7, A and B). Interestingly, we observed longer calcium transients in Ctl cells compared to GluN1KO cells (Fig. S7C), which we binned into time intervals for further investigation. While the quantity of short events ranging from 0 – 5 s did not significantly differ, Ctl cells exhibited a greater number of events ranging from 5 – 25 s and 25 – 150 s. Strikingly, 8 out of 18 Ctl cells (44%) had a calcium transient lasting between 150 – 350 s in duration, in contrast to 0 out of 15 GluN1KO cells (Fig. 4F). We examined the kernel density estimation (KDE) of Ctl and GluN1KO datapoints within the intervals to compare their underlying distribution. Whereas probability density functions overlapped between 0 – 5 s, Ctl functions exhibited a rightward shift along the x-axis during longer intervals (Fig. 4G), suggesting NMDAR involvement in long calcium transients. To integrate space, time and intensity, we summed all ΔF/F values within events, revealing stronger transients in Ctl cells (Fig. S7D). Binning values into different ranges, we noticed that Ctl cells had a higher amount of small and large events compared to GluN1KO cells (Fig. 4H). The larger events seemed primarily driven by the long transients, as visualized in the nonbinary raster plots (Fig. 4, C and D). These data indicate that NMDARs are frequently activated early in development and give rise to robust and sustained calcium transients which may influence synapse formation and dendritic maturation.

**Fig. 4.**
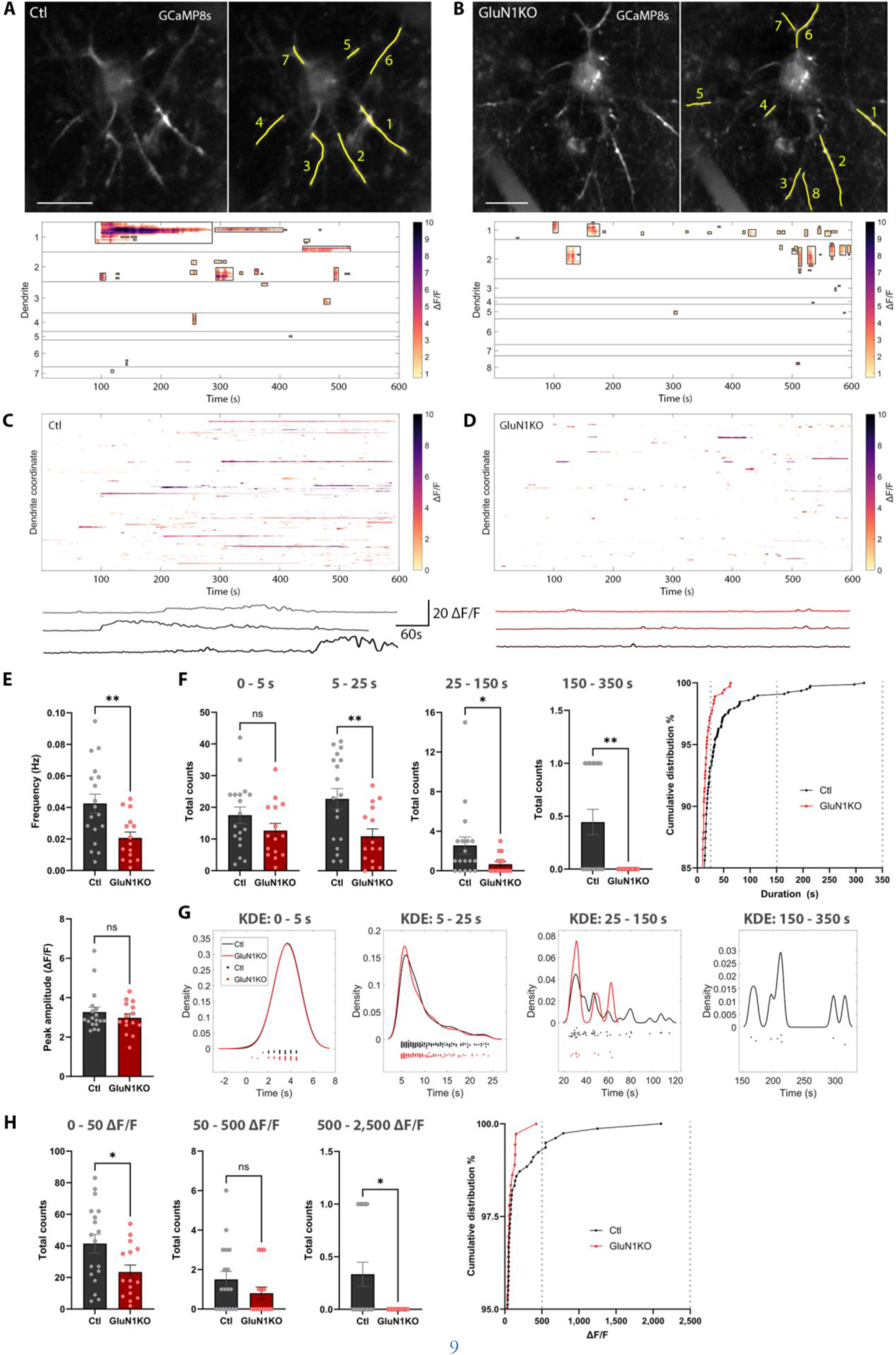
NMDARs are frequently activated during development and give rise to long calcium transients. (**A, B**) 2-photon maximum projection images of an example Ctl and GluN1KO neuron and 2D-reconstructions of dendrites. The raster plots depict changes in fluorescence (ΔF/F) at coordinates along concatenated dendrites from each neuron. Events are bordered by black bounding boxes. Scale bar, 5 μm. (**C, D**) Raster plots displaying events of Ctl cells (**C**), and GluN1KO cells (**D**) (1,826 Ctl dendrite coordinates, 18 cells from 3 mice; 1,765 GluN1KO dendrite coordinates, 15 cells from 2 mice). Exemplary ΔF/F traces from 3 Ctl (**C**) and GluN1KO (**D**) cells. (**E**) Graphs of frequency, normalized by dendritic length, and amplitude measurements (n = 18 Ctl cells from 3 mice; n = 15 GluN1KO cells from 2 mice). \*\**P* = 0.0059 (unpaired *t* test) and *P* = 0.5800 (Mann-Whitney test). (**F**) Event counts over 10 minute imagine period, binned into different durations and cumulative distribution of durations (n = 18 Ctl cells from 3 mice; n = 15 GluN1KO cells from 2 mice). 0 - 5 s: *P* = 0.1776 (unpaired *t* test). 5 - 25 s: \*\**P* = 0.0096 (Mann-Whitney test). 25 - 150 s: \**P* = 0.0339 (Mann-Whitney test). 150 - 350 s: \*\**P* = 0.0036 (Mann-Whitney test). (**G**) Kernel density estimation of Ctl and GluN1KO data within individual time intervals. (**H**) Sum of ΔF/F values within events binned into different ranges and cumulative distribution of ΔF/F sums (n = 18 Ctl cells from 3 mice; n = 15 GluN1KO cells from 2 mice). 0 - 50 ΔF/F: \**P* = 0.0244 (unpaired *t* test). 50 - 500 ΔF/F: *P* = 0.2077 (Mann-Whitney test). 500 - 2500 ΔF/F: \**P* = 0.0213 (Mann-Whitney test).

### Structural changes following long calcium transients

We next investigated whether the robust NMDAR-mediated calcium transients were associated with structural changes in the dendritic arbor. Upon closer examination, we indeed identified structural changes occurring with long calcium events (Fig. 5A, Fig. S8, A and B). Following the initiation of the calcium transient in the parent dendrite, we observed filopodia branching at the site of activation on the timescale of seconds (Movie S1-3). Initially, we attempted to simultaneously image GCaMP8s and tdTomato to capture calcium transient responses and structural changes separately. Unfortunately, cells expressing both GCaMP8s and tdTomato showed very few calcium responses, likely due to intramolecular interference (Fig. S10, A-D). To confirm that we were observing structural changes, we analysed pixel intensities of lines drawn along the perimeter of Ctl parent dendrites before and after the calcium transient. The pixel intensities revealed intensity peaks present only after the calcium transient, resembling intersections of the lines with newly grown filopodia (Fig. 5B, Fig. S8, C and D). Convincingly, preexisting branching filopodia can be seen at the onset of the calcium transient, indicating a widespread activation of nearby filopodia (Fig. S8, C and D). Moreover, short calcium transients in Ctl dendrites were not linked with structural changes (Fig. 5C). In contrast, GluN1KO cells did not exhibit any structural changes linked to calcium transients, even in dendrites displaying frequent events (Fig. 5D, Fig. S9, Movie S4). In summary, these data provide evidence that sustained calcium signalling linked to NMDAR activation can contribute to filopodial growth, reinforcing our previous findings.

**Fig. 5.**
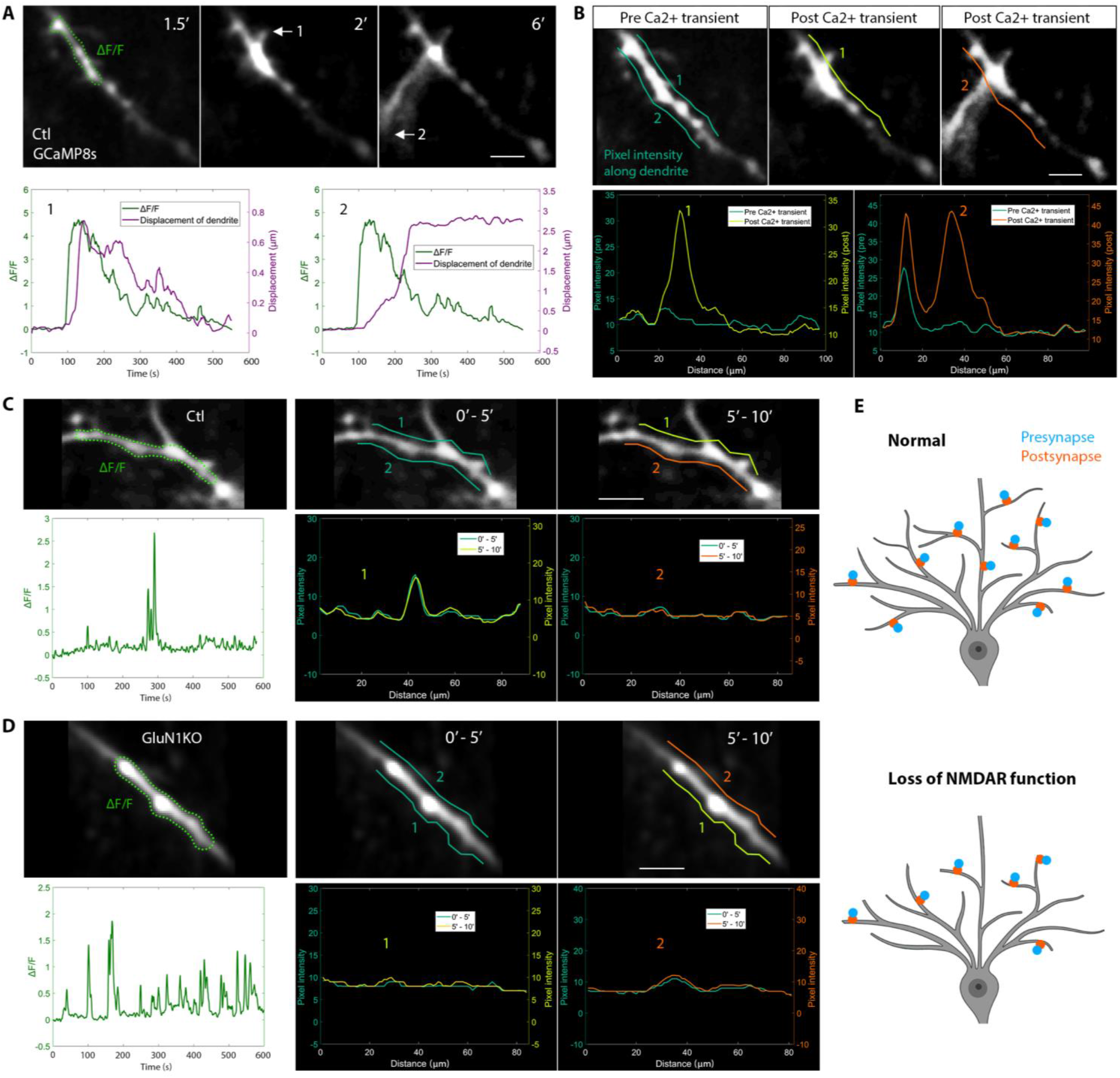
NMDAR-mediated long calcium transients are coupled to local structural changes in dendritic arbor. (**A**) Time-lapse images of a GCaMP8s+ Ctl cell. The ΔF/F ROI is outlined by the dashed green line. Two branching filopodia are denoted by white arrows. The lower panels show mean fluorescence activity (green) and dendritic displacement (purple) traces for each filopodial branch, providing a visual representation of transient onset and dendritic branching. Scale bar, 1 μm. (**B**) Assessment of filopodial growth by line intensity scanning alongside dendrites before and after calcium transient. Lines prior to calcium transient are presented in cyan and superimposed onto images after the calcium transient. Pixel intensity traces before (cyan) and after (yellow / orange) the calcium transient are overlaid, illustrating changes in fluorescence. Scale bar, 1 μm. (**C**) Line intensity scanning alongside a Ctl dendrite to assess whether structural changes occur following short calcium transients. Scale bar, 1 μm. (**D**) Line intensity scanning alongside a highly active GluN1KO dendrite to assess whether structural changes have occurred. Scale bar, 1 μm. (**E**) Final schematic illustrating basal dendrites of CA1 PNs, summarizing findings. Loss of NMDAR function results in a reduction in synapse densities and dendritic complexity.

## Discussion

Our experiments demonstrate a role for NMDARs in shaping neuronal connections in the hippocampus, which is dependent on the timing of signaling and the anatomical circuit studied. We found a decrease in synapse densities specifically for SC connections by blocking NMDAR function, both *in vitro* and *in vivo*, alongside reductions in dendritic arbor complexity (Fig. 5E).

Our results highlight the application of functional *in vivo* imaging at these very early stages, with the surprising finding of minute-long, dendritic calcium transients that immediately preceded the growth of filopodia and were completely lost in the absence of NMDAR function. NMDAR-independent, local dendritic calcium events lasting < 1 second and appearing after presynaptic contact have been observed in developing hippocampal neurons *in vitro* (*14*), and indeed we see similar brief, non-NMDAR-dependent events occurring *in vivo*. We previously identified spontaneous NMDAR-dependent calcium transients in discrete sections of dendrites in dissociated hippocampal neurons, which, while longer, only lasted up to 10 seconds (*8*). We are not aware of such prolonged dendritic events being previously described and it will be interesting in future to explore their nature in more detail. The association of these NMDAR-mediated transients with structural changes provides a potential cellular underpinning for the alterations in dendritic branching and synapse densities seen following deletion of NMDAR function. Interestingly, pharmacological blockade of glutamatergic signalling, including NMDARs, restricts both filopodial motility and dendritic branch development in retinal ganglion cells and *Xenopus* tectal neurons (*10, 15*). While *in vitro* studies of mammalian cortical and hippocampal pyramidal neurons found no effects from similar manipulations (*16*), these were carried out at a later developmental stage. Therefore, our results suggest that the importance of NMDARs is dependent on developmental timing rather than playing no role in mammalian pyramidal neurons. Our data provide *in vivo* evidence in mammalian central nervous system neurons that supports the link between NMDAR function, filopodial growth, dendritic arbor shaping and synaptogenesis (*17-19*).

Our results demonstrating the absence of any effects of blocking action potential firing support research indicating minimal involvement of evoked transmitter release in synapse formation (*12, 20, 21*) and filopodial motility (*15*). Given our previous work indicating that stochastic, spontaneous release occurs at exceptionally high levels in young cultured hippocampal neurons (*22*), this mode of release may dominate during our timing of intervention (E14 – P3). Moreover, early spontaneous release has been shown to activate distant dendritic NMDARs *in vitro* (*8*). Therefore it is possible that stochastic release of glutamate from developing SC axons could activate CA1 dendritic NMDARs. Interestingly, specific pools of NMDARs on the postsynaptic membrane are known to be targeted by spontaneous release (*23*). However, further investigation is needed to understand the precise source of glutamate.

Our findings on the timing of intervention align with seminal research showing that the postnatal blockade of NMDAR function does not affect synaptic or dendritic structure (*11*). Having obtained similar results in postnatal organotypic cultures, we targeted our genetic manipulations to embryonic stages, thereby uncovering structural and functional impacts upon NMDAR silencing in an early developmental window. Notably, we found no effect of abolishing NMDAR function in this window on synaptic connections from EC inputs, indicating some degree of specificity. We hypothesize that this lack of phenotype in the distal tuft dendrites is likely to be due to the timing of their formation. We found that synapses were already present in the distal tufts at P0 and did not change in density between then and P3, in striking contrast to SC inputs. Projections from the EC are already present in this region, the future stratum lacunosum moleculare, as early as E17 (*24*), making it likely that EC inputs form during embryonic development. It remains possible that NMDARs could still play a role in the early formation of these inputs if this predates the abolition of NMDAR function in our experiments. While some synapses still form in the absence of NMDAR function, we attribute a time-sensitive and synapse-specific role to NMDARs during circuit and dendrite formation which could allow for optimization of hippocampal connectivity. Given the strong association for mutations in NMDAR subunit genes with neurodevelopmental disorders (*25*), this could also provide a mechanistic link between loss of NMDAR function and cognitive impairment seen in these disorders.

## Author contributions

Conceptualization: L.C.A. Methodology: L.C.A, N.S.L, N.F.H., S.K., F.D. and A.K. Investigation and analysis: N.S.L. Funding acquisition: L.C.A, F.D. and A.K. Project administration: L.C.A. Supervision: L.C.A., N.F.H., S.K., F.D. and A.K. Writing – original draft: N.S.L. and L.C.A. with input from all authors.

## Data and materials availability

All data are available in the main text or the supplementary materials. The code developed for analysis is available on GitHub [https://github.com/lcandreae/Andreae-Lab/blob/main/CalciumImaging_neuronJdendrites.m]

## Supplementary Materials

Materials and Methods

Figs. S1 to S10

Movies S1 to S4

## References

1. L. C. Katz, C. J. Shatz, Synaptic activity and the construction of cortical circuits. Science 274, 1133–1138 (1996).

2. C. L. Torborg, M. B. Feller, Spontaneous patterned retinal activity and the refinement of retinal projections. Prog. Neurobiol. 76, 213–235 (2005).

3. J. R. Sanes, J. W. Lichtman, Development of the vertebrate neuromuscular junction. Annu. Rev. Neurosci. 22, 389–442 (1999).

4. A. Bleckert, R. O. Wong, Identifying roles for neurotransmission in circuit assembly: insights gained from multiple model systems and experimental approaches. BioEssays 33, 61–72 (2011).

5. D. Kerschensteiner, J. L. Morgan, E. D. Parker, R. M. Lewis, R. O. Wong, Neurotransmission selectively regulates synapse formation in parallel circuits in vivo. Nature 460, 1016–1020 (2009).

6. G. Y. Wu, R. Malinow, H. T. Cline, Maturation of a central glutamatergic synapse. Science 274, 972–976 (1996).

7. T. V. Bliss, G. L. Collingridge, A synaptic model of memory: long-term potentiation in the hippocampus. Nature 361, 31–39 (1993).

8. L. C. Andreae, J. Burrone, Spontaneous neurotransmitter release shapes dendritic arbors via long-range activation of NMDA receptors. Cell Reports 10, 873–882 (2015).

9. H. Mizuno, W. Luo, E. Tarusawa, Y. M. Saito, T. Sato, Y. Yoshimura, S. Itohara, T. Iwasato, NMDAR-regulated dynamics of layer 4 neuronal dendrites during thalamocortical reorganization in neonates. Neuron 82, 365–379 (2014).

10. W. C. Sin, K. Haas, E. S. Ruthazer, H. Cline, Dendrite growth increased by visual activity requires NMDA receptor and Rho GTPases. Nature 419, 475–480 (2002).

11. W. Lu, E. A. Bushong, T. P. Shih, M. H. Ellisman, R. A. Nicoll, The cell-autonomous role of excitatory synaptic transmission in the regulation of neuronal structure and function. Neuron 78, 433–439 (2013).

12. D. L. Barabási, G. F. P. Schuhknecht, F. Engert, Functional neuronal circuits emerge in the absence of developmental activity. Nat. Commun. 15, 364 (2024).

13. L. Luo, Architectures of neuronal circuits. Science 373, eabg7285 (2021).

14. C. Lohmann, T. Bonhoeffer, A role for local calcium signaling in rapid synaptic partner selection by dendritic filopodia. Neuron 59, 253–260 (2008).

15. W. T. Wong, R. O. Wong, Changing specificity of neurotransmitter regulation of rapid dendritic remodeling during synaptogenesis. Nat. Neurosci. 4, 351–352 (2001).

16. A. Dunaevsky, A. Tashiro, A. Majewska, C. Mason, R. Yuste, Developmental regulation of spine motility in the mammalian central nervous system. Proc. Natl Acad. Sci. USA 96, 13438–13443 (1999).

17. H. T. Cline, K. Haas, The regulation of dendritic arbor development and plasticity by glutamatergic synaptic input: a review of the synaptotrophic hypothesis. J. Physiol. 586, 1509–1517 (2008).

18. J. E. Vaughn, Fine structure of synaptogenesis in the vertebrate central nervous system. Synapse 3, 255–285 (1989).

19. C. M. Niell, M. P. Meyer, S. J. Smith, In vivo imaging of synapse formation on a growing dendritic arbor. Nat. Neurosci. 7, 254–260 (2004).

20. C. W. Lin, S. Sim, A. Ainsworth, M. Okada, W. Kelsch, C. Lois, Genetically increased cell-intrinsic excitability enhances neuronal integration into adult brain circuits. Neuron 65, 32–39 (2010).

21. Z. Molnár, G. López-Bendito, J. Small, L. D. Partridge, C. Blakemore, M. C. Wilson, Normal development of embryonic thalamocortical connectivity in the absence of evoked synaptic activity. J. Neurosci. 22, 10313–10323 (2002).

22. L. C. Andreae, N. B. Fredj, J. Burrone, Independent vesicle pools underlie different modes of release during neuronal development. J. Neurosci. 32, 1867–1874 (2012).

23. A. L. Reese, E. T. Kavalali, Single synapse evaluation of the postsynaptic NMDA receptors targeted by evoked and spontaneous neurotransmission. eLife 5, e21170 (2016).

24. H. Supèr, E. Soriano, The organization of the embryonic and early postnatal murine hippocampus. II. Development of entorhinal, commissural, and septal connections studied with the lipophilic tracer DiI. J. Comp. Neurol. 344, 101–120 (1994).

25. A. García-Recio, A. Santos-Gómez, D. Soto, N. Julia-Palacios, À. García-Cazorla, X. Altafaj, M. Olivella, GRIN database: A unified and manually curated repertoire of GRIN variants. Hum. Mutat. 42, 8–18 (2021).

26. L. Stoppini, P. A. Buchs, D. Muller, A simple method for organotypic cultures of nervous tissue. J. Neurosci. Methods 37, 173–182 (1991).

27. R. A. Ellingford, M. J. Panasiuk, E. R. de Meritens, R. Shaunak, L. Naybour, L. Browne, M. A. Basson, L. C. Andreae, Cell-type-specific synaptic imbalance and disrupted homeostatic plasticity in cortical circuits of ASD-associated Chd8 haploinsufficient mice. Mol. Psychiatry 26, 3614–3624 (2021).

28. P. C. Lee, H. Y. He, C. Y. Lin, Y. T. Ching, H. T. Cline, Computer aided alignment and quantitative 4D structural plasticity analysis of neurons. Neuroinformatics 11, 249–257 (2013).

29. M. Pachitariu, C. Stringer, M. Dipoppa, S. Schröder, L. F. Rossi, H. Dalgleish, M. Carandini, K. D. Harris, Suite2p: beyond 10,000 neurons with standard two-photon microscopy. bioRxiv 061507 (2017).

30. E. Meijering, M. Jacob, J. C. F. Sarria, P. Steiner, H. Hirling, M. Unser, Design and validation of a tool for neurite tracing and analysis in fluorescence microscopy images. Cytometry A. 58, 167–176 (2004).

