## Supplemental Files for "NMDA receptor activation drives early synapse formation *in vivo*"

**The file includes:**

Materials and Methods
Figs. S1 to S10

**Other Supplementary Materials for this manuscript include the following:**

Movies S1 to S4

### Materials and Methods

Animals

All procedures were performed according to the Animals (Scientific Procedures) Act 1986 with ethical approval granted by the UK Home Office. *Grin1*$^{fl/fl}$ and WT C57BL/6 mice between E18.5 and 5 months of age were used for all experiments. Animals were kept in a temperature- and humidity-regulated room under a 12-hour light cycle, with access to food and water at all times. Animal health and welfare was monitored daily by technicians supervised by a designated veterinarian. Sex was not considered during data acquisition and analysis.

Hippocampal organotypic cultures

Brains were placed in filtered dissecting solution upon decapitation and hippocampi dissected out. 300 µm organotypic slices were prepared and cultured on Millicell membranes (*26*), previously coated in poly-L-lysine (100µg/ml) and laminin (100µg/ml) overnight in a humidified incubator (37°C, 5% CO2). Four hours after culturing, a 20 µL droplet of AAV9-hSyn-eGFP was added onto each slice. The culture medium was replaced at 1DIV with medium containing pharmacological blockers (100 µM APV or 1 µM TTX). Slices were maintained in a humidified incubator for 48 hr before fixation.

Histology on organotypic cultures

At 3DIV hippocampal organotypic slices were fixed in 4% PFA and 4% sucrose in PBS for 30 min. Slices were permeabilized for 30 min (0.5% Triton X-100 in PBS) and maintained in blocking buffer for 3 hr (0.3% Triton X-100, 5% BSA, 0.2 M glycine in PBS). Organotypic slices were incubated in primary antibodies specific for GFP (GFP; 1:1000, chicken anti-GFP, Abcam), homer (homer; 1:1000, rabbit anti-homer, Synaptic Systems) and bassoon (bassoon; 1:1000, mouse anti-bassoon) diluted in blocking buffer for 48 hr at 4°C. Slices were washed with blocking buffer and incubated in secondary antibodies (goat α-chicken Fluor488, goat α-rabbit Fluor555 and goat α-mouse IgG2a Fluor647, 1:1000, Alexa) diluted in blocking buffer for further 48 hr at 4°C. Sections were mounted onto glass slides with Mowiol (Sigma).

*In utero* virus injections

Prior to the procedure, pregnant dams at day E14.5 of gestation received a subcutaneous injection of Vetergesic (Ceva Animal Health) at a dose of 0.1 mg/kg. Anaesthesia was maintained at 2% isoflurane during surgery. A midline incision was made and embryos were carefully extracted from the abdomen. Embryos were kept moist with sterile PBS warmed to 32°C throughout the procedure. A virus solution with 0.005% Fast Green dye was injected into a single lateral ventricle of embryos. Following closure of the abdominal wall, dams recovered for 1 hr in a 35°C recovery chamber and returned to their cages with moistened food.

Whole-cell patch clamp electrophysiology

Acute brain slices of 300 µm thickness were obtained as previously described (*27*). Slices recovered in magnesium-free artificial cerebrospinal fluid (ACSF; 124 mM NaCl, 5 mM KCl, 1.25 mM $Na_2HPO_4$, 2 mM $CaCl_2$, 26 mM $NaHCO_3$ and 20 mM D-glucose) equilibrated with 95% O2/5% CO2 for 45 min at 34°C. During recordings slices were held at room temperature in ACSF containing 1 µM TTX, 20 µM NBQX and 30 µM gabazine (HEKA EPC10/2 amplifier, Pulse software). Borosilicate glass electrodes of 3 - 7 MΩ resistance were filled with K-gluconate internal solution (135 mM K-gluconate, 10 mM KCl, 10 mM HEPES, 1 mM MgCl2, 2 mM Na-adenosine triphosphate ($Na_2ATP$) and 0.4 mM Na-guanosine triphosphate ($Na_3GTP$)). tdTomato+ cells in CA1 were voltage clamped at -60 mV and NMDAR-mediated currents recorded following a localized picospritzer puff of 100 µM NMDA and 10 µM glycine for 100 ms.

Perfusions and histology at P0 and P3

Pups were transcardially perfused with 2 ml PBS followed by 15 ml 4% PFA in PBS. Brains were held in 4% PFA for 24 hours at 4°C following perfusion fixation. To prepare brain slices, brains were embedded in 2% agarose (Thermo Scientific) and 150 µm coronal slices obtained using a Leica VT 1000S vibratome. Sections were permeabilized for 30 min (0.5% Triton X-100 in PBS) and placed in blocking buffer for 3 hr (0.3% Triton X-100, 3% BSA, 10% Goat Serum, 0.02% NaN3, 0.2M glycine in PBS). Slices were incubated in primary antibodies specific for GFP (GFP; 1:1000, chicken anti-GFP, Abcam), homer (homer; 1:1000, rabbit anti-homer, Synaptic Systems) and bassoon (bassoon; 1:1000, mouse anti-bassoon, Abcam) diluted in blocking buffer for 48 hr at 4°C. Slices were washed with blocking buffer and incubated overnight at 4°C in secondary antibodies (goat α-chicken Fluor488, goat α-rabbit Fluor555 and goat α-mouse IgG2a Fluor647, 1:1000, Alexa) diluted in blocking buffer. Brain sections were mounted onto glass slides with Mowiol.

Histology image acquisition & analysis

Z-stacks of dendrites and synapses from GFP+ CA1 PNs were imaged using a Zeiss LSM 800 confocal microscope under a 63x objective. Dendrites were reconstructed using the filament tracer function in IMARIS (BitPlane) from which dendritic length was quantified. The Spots function was used to detect homer puncta inside the GFP-labelled dendrite. The homer puncta were subsequently tested for colocalization with presynaptic bassoon puncta to detect structural synapses. Homer and synapse quantities were normalized by dendritic length to obtain density measurements. To acquire images of whole-cell morphology, z-stacks were taken under a 20x objective. The filament tracer tool in IMARIS was used to reconstruct the entire neuronal arbor.

Picospritzer activation of NMDARs in SO, SR and SLM

Acute brain slices were transferred to an imaging chamber superfused with oxygenated (95% $O_2$/5% $CO_2$) ACSF at room temperature. Picospritzer pipettes filled with 5 µM Alexa Fluor 594, 100 µM NMDA and 10 µM glycine were localized to mRuby+ dendritic regions in CA1 guided by 2-photon excitation with a Ti-Sapphire laser (Coherent Chameleon) at 1060 nm wavelength. Photon emission was collected by a GaAsP PMT using a 40x objective (Olympus, 0.8 NA) upon GCaMP6s excitation at 920 nm in response to 200 ms picospritzer puffs. Images were collected at 30 Hz with a rolling average of 25 frames at a resolution of 512 x 512 pixels. Upon background subtraction, changes in GCaMP6s fluorescence ($\Delta F/F$) relative to baseline signals were measured within dendritic ROIs.

Cranial window implant

Prior to the implant surgery, a 3 mm diameter circular glass coverslip (Generon) was glued onto a brass metal cannula of 3 mm diameter using UV-curable glue (NOA 81, Norland Optical Adhesives). Mice were given a subcutaneous injection of Metacam (20 mg/kg), dexamethasone (16 mg/kg), and atropine (0.41 mg/kg).

Anaesthesia was maintained between 1.5% and 2.5% during surgery. Body temperature was controlled by placing mice on a heating mat set to 38°C. The left skull hemisphere was covered with a layer of cyanoacrylate glue and a headplate secured in place with UV-curable cement (Venus Diamond Flow, Kulzer). The coordinates of the window implant in the right hemisphere were visually approximated. A craniotomy was performed, and cortical tissue gently aspirated under constant saline irrigation to expose the dorsal hippocampus. The hippocampus was covered with a thin layer of Kwik-Sil (WPI) before inserting the cannula above the hippocampus. UV-curable cement was applied to affix the edges of the cannula to the skull. A subcutaneous injection of saline and methadone (0.08 mg/kg) was administered postoperatively. Mice were placed in a heating chamber to recover before imaging.

*In vivo* 2-photon imaging & analysis

*In vivo* 2-photon imaging experiments were conducted following cranial implant surgery. Anaesthesia was maintained between 0.6 - 1% during imaging. In separate experiments, GCaMP8s and tdTomato were excited with a Ti-Sapphire laser (Coherent Chameleon) at 920 nm and 1040 nm wavelength, respectively. Emitted light was collected by a GaAsP PMT using a 16x objective (Nikon, 0.8 NA).

For continuous structural imaging of tdTomato+ dendrites, a single plane was imaged at 15 Hz for 10 min using a rolling average of 100 frames at 1024 x 1024 pixels. In-plane distal dendrites were imaged towards the periphery of dendritic arbors, while in-plane proximal dendrites were imaged near the soma (such that the soma can be visualized in 7 of 9 cells). Time-series were pre-processed in ImageJ using the SIFT plugin for motion correction, followed by the use of a 3D median filter (2 x 2 x 2, in x-y-z). Filopodial tips branching from parent dendrites were tracked across frames using the Spots function in IMARIS (Bitplane). Their displacement was computed by determining the distance of each coordinate along the track from the initial coordinate at imaging onset. The distance covered represents the length of the entire tracked path.

For structural imaging of tdTomato+ CA1 basal arbors, z-stacks of 1 µm depth were acquired at 15 Hz. At each depth 100 frames were averaged to obtain high resolution images at 1024 x 1024 pixels. Z-stacks were pre-processed in ImageJ by applying a 3D median filter (2 x 2 x 2, in x-y-z) to each image. Basal arbors at timepoints 0 hr and 2 hr were reconstructed using the filament tracer tool within IMARIS. Dendrites were subsequently aligned using 4DSPA software to identify retracted, added and stable dendrites (*28*).

2-photon calcium imaging experiments were performed at 30 Hz for 10 min with a rolling average of 15 frames at a resolution of 512 x 512 pixels. Time-series were pre-processed in Suite2p for motion correction (*29*), and ImageJ for application of a 3D median filter (2 x 2 x 2, in x-y-z). Frames containing action potentials were removed to restrict analysis to local calcium transients. Dendrites were traced in NeuronJ (*30*) and changes in fluorescence ($\Delta F/F$) at dendritic coordinates were computed using custom-written MATLAB code for automated detection of events. In summary, the baseline for ΔF/F computation was determined by calculating the mean of the lowest 10% values in each trace. Traces were subsequently smoothed, and values exceeding 30 times the standard deviation of the individual noise band were accepted for event detection. Raster plots were analysed to detect events connected in time (across frames) and space (across neighbouring coordinates). Events smaller than 6 pixels in size were defined as noise and excluded from the analysis.

Structure-function analysis

Structure-function analysis was performed on calcium imaging time-series. For functional analysis, ROIs were drawn around segments of parent dendrites and changes in mean fluorescence ($\Delta F/F$) were computed across all frames. For structural analysis, filopodial tips were tracked across frames using the Spots function in IMARIS to obtain displacement information. Structure and function traces were overlaid to visualize the onset of the calcium transient and filopodial growth.

To test whether structural changes have occurred, all images taken before and after the calcium transient were averaged to obtain two images – 'Pre Ca2+ transient' & 'Post Ca2+ transient'. Image contrasts were enhanced to reveal dim structures. Lines were drawn beside dendrites in images before the calcium transient and overlaid onto respective images after the calcium transient. Pixel intensities were measured along the drawn lines before and after the calcium transient, and traces were superimposed. Intensity peaks that are present after the calcium transient in Ctl cells represent intersections of the lines with newly formed filopodia. Since GluN1KO cells exhibited no structural changes associated with calcium transients, we analysed averaged images from the first and second halves of the calcium imaging time-series ('Pre Ca2+ transient' = 0 – 5 min; 'Post Ca2+ transient' = 5 – 10 min).

Statistics

All statistical analyses were performed using GraphPad Prism. Data are presented as mean ± standard error of the mean. For pairwise and multiple comparisons the D'Agostino-Pearson normality test was used to assess Gaussian distribution within each data set. Comparisons of normally distributed datasets were either performed using unpaired *t* tests, paired *t* tests, or one-way ANOVAs followed by Tukey's multiple comparisons tests. Comparisons of non-normal data sets were either performed using nonparametric Mann-Whitney tests, Wilcoxon matched-pairs signed rank tests or Kruskal Wallis tests followed by Dunn's multiple comparisons tests. *P*-values < 0.05 were accepted as statistically significant: *$P < 0.05$, **$P < 0.01$, ***$P < 0.001$ and ****$P < 0.0001$.

**Fig. S1.**

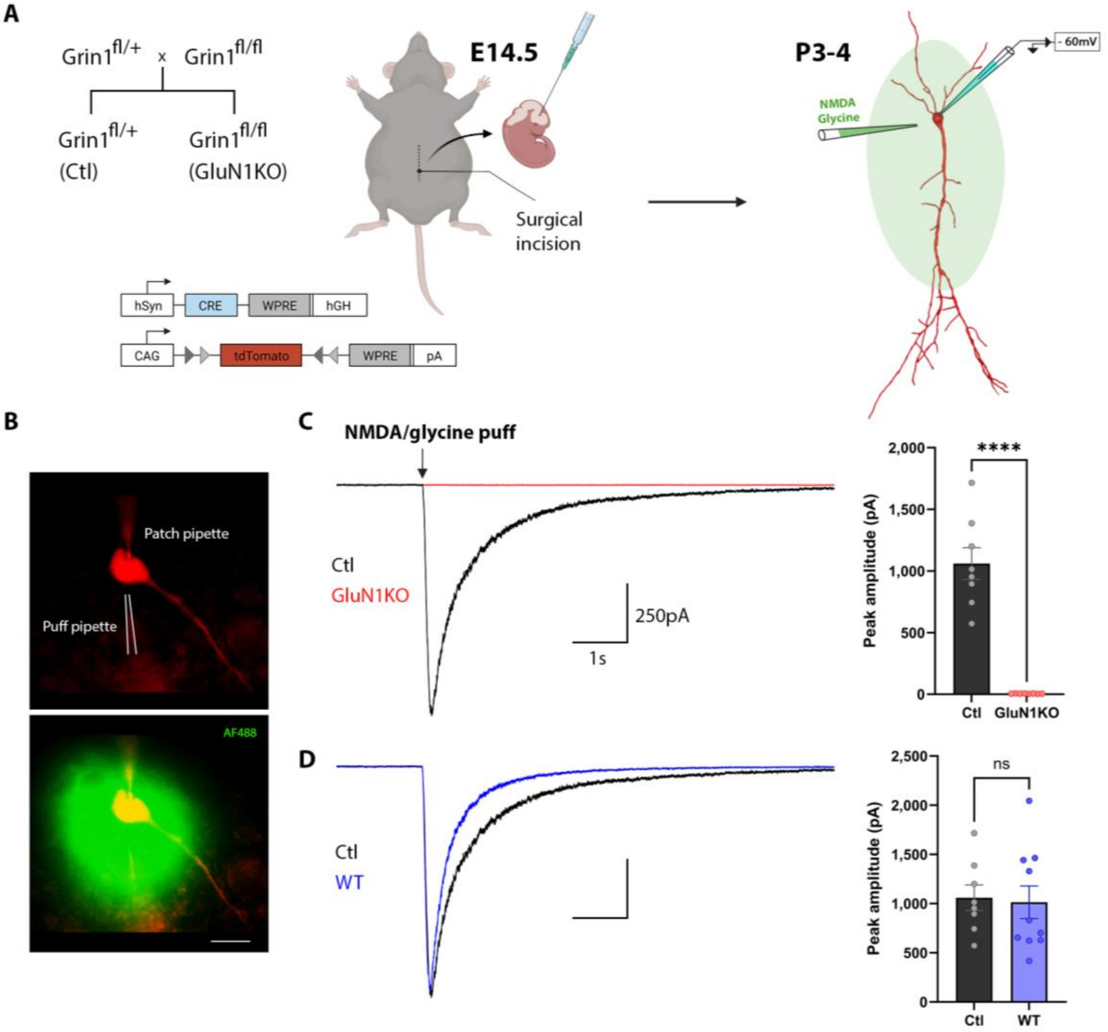


**Fig. S1. Loss of NMDAR function in CA1 PNs at P3-4 by administering AAV-Cre during embryonic development.** (**A**) Breeding strategy; schematic of E14.5 *in utero* viral injection of AAV-Cre and AAV-flex-tdTomato; whole-cell patch clamp recordings of CA1 PNs at P3-4, targeted by focal picospritzer puffs of NMDA and glycine. (**B**) Visualization of a picospritzer puff of AF488 onto a tdTomato+ neuron. Scale bar, 10 µm. (**C**) Mean current traces and peak amplitude responses to NMDA and glycine application (n = 8 Ctl cells from 2 mice; n = 8 GluN1KO cells from 3 mice). ****$P < 0.0001$ (unpaired *t* test). (**D**) Comparison of mean somatic current traces and peak amplitudes from Ctl and wildtype (WT) cells (n = 8 Ctl cells from 2 mice; n = 10 WT cells from 2 mice). $P$ = 0.8348 (unpaired *t* test).

**Fig. S2.**

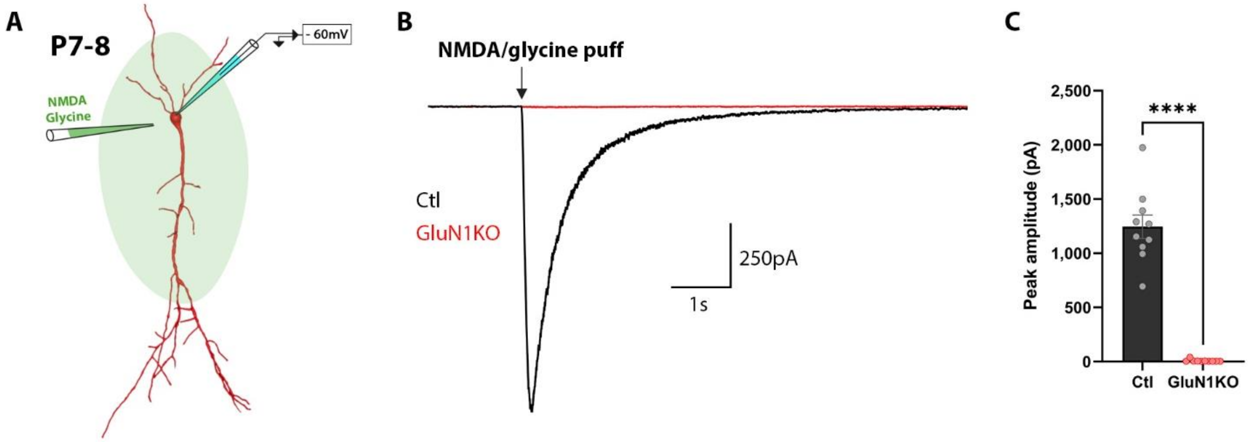


**Fig. S2. Sustained loss of NMDAR function in CA1 PNs at P7-8.**
(**A**) Schematic representation of a whole-cell patch clamped neuron at P7-8, targeted by a picospritzer puff of NMDA and glycine. (**B**) Population mean current trace of Ctl and GluN1KO neurons. (**C**) Comparison of peak amplitudes from Ctl and GluN1KO cells (n = 10 Ctl cells from 2 mice; n = 10 GluN1KO cells from 3 mice). ****$P < 0.0001$ (Mann-Whitney test).

**Fig. S3.**

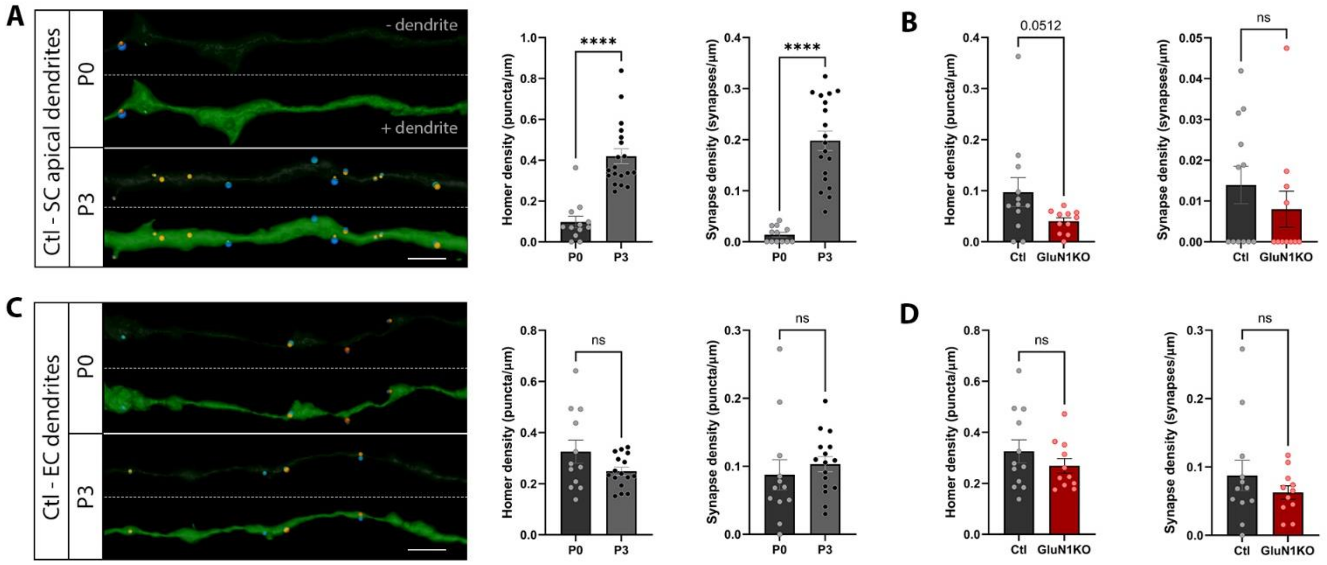


**Fig. S3. Unique timing of synapse formation of SC and EC inputs at dendrites in CA1.** (**A**) Reconstructions of SC apical dendrites and synapses of Ctl neurons at P0 and P3, along with quantification of homer and synapse densities (n = 12 Ctl cells from 3 P0 mice; n = 19 Ctl cells from 5 P3 mice). Homer density: ****$P < 0.0001$ (Mann-Whitney test). Synapse density: ****$P < 0.0001$ (unpaired *t* test). Scale bar, 5 µm. (**B**) Comparison of homer and synapse densities in SC dendrites between Ctl and GluN1KO cells at P0 (n = 12 Ctl cells from 3 mice; n = 11 GluN1KO cells from 3 mice). Homer density: $P = 0.0512$. Synapse density: $P = 0.6947$ (Mann-Whitney tests). (**C**) Reconstructions of apical tuft dendrites and synapses of Ctl neurons at P0 and P3, and quantification of homer and synapse densities (n = 12 Ctl cells from 3 P0 mice; n = 16 Ctl cells from 4 P3 mice). Homer density: $P = 0.0839$ (unpaired *t* test). Synapse density: $P = 0.1627$ (Mann-Whitney test). Scale bar, 5 µm. (**D**) Comparison of homer and synapse densities in EC dendrites between Ctl and GluN1KO cells at P0 (n = 12 Ctl cells from 3 mice; n = 11 GluN1KO cells from 3 mice). Homer density: $P = 0.3098$ (unpaired *t* test). Synapse density: $P = 0.6629$ (Mann-Whitney test).

**Fig. S4.**

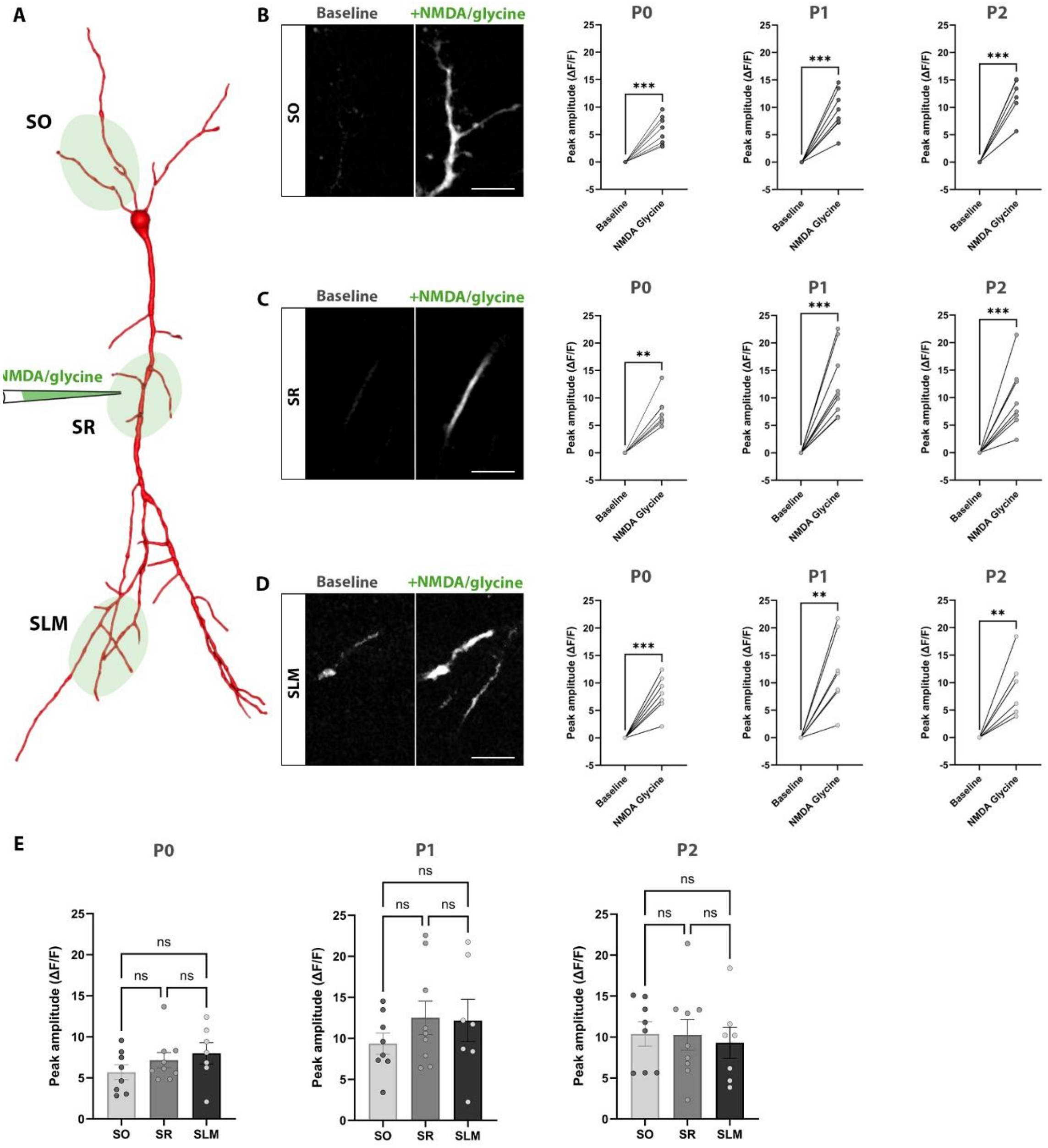


**Fig. S4. NMDARs are expressed and functional during SC synapse formation.**
(**A**) Schematic showing 2-photon guided localized picospritzer puffs to SO, SR and SLM dendrites expressing mRuby2 and GCaMP6s. (**B-D**) Images depicting baseline GCaMP6s signal and increase in fluorescence in dendrites following NMDA and glycine administration. Scale bar, 10 µm. (**B**) Quantification of peak amplitudes in SO at P0 (n = 8 cells from 2 WT mice; ***$P = 0.0004$, paired *t* test), P1 (n = 8 cells from 2 WT mice; ***$P = 0.0002$, paired *t* test), and P2 (n = 8 cells from 2 WT mice; ***$P = 0.0002$, paired *t* test). (**C**) Peak amplitudes in SR at P0 (n = 9 cells from 2 WT mice; **$P = 0.0039$, Wilcoxon test), P1 (n = 9 cells from 2 WT mice; ***$P = 0.0003$, paired *t* test), and P2 (n = 9 cells from 2 WT mice; ***$P = 0.0006$, paired *t* test). (**D**) Peak amplitudes in SLM at P0 (n = 7 cells from 2 WT mice; ***$P = 0.0008$, paired *t* test), P1

(n = 7 cells from 2 WT mice; $**P = 0.0033$, paired *t* test), and P2 (n = 7 cells from 2 WT mice; $**P = 0.0027$, paired *t* test). (**E**) Comparison of peak amplitudes between dendritic compartments at P0 ($P > 0.9999$ for SO versus SR, $P = 0.4501$ for SO versus SLM, and $P > 0.9999$ for SR versus SLM; Kruskal-Wallis test), P1 ($P = 0.5010$ for SO versus SR, $P = 0.6107$ for SO versus SLM, and $P = 0.9927$ for SR versus SLM; one-way ANOVA), and P2 ($P = 0.9992$ for SO versus SR, $P = 0.9120$ for SO versus SLM, and $P = 0.9218$ for SR versus SLM; one-way ANOVA).

**Fig. S5.**

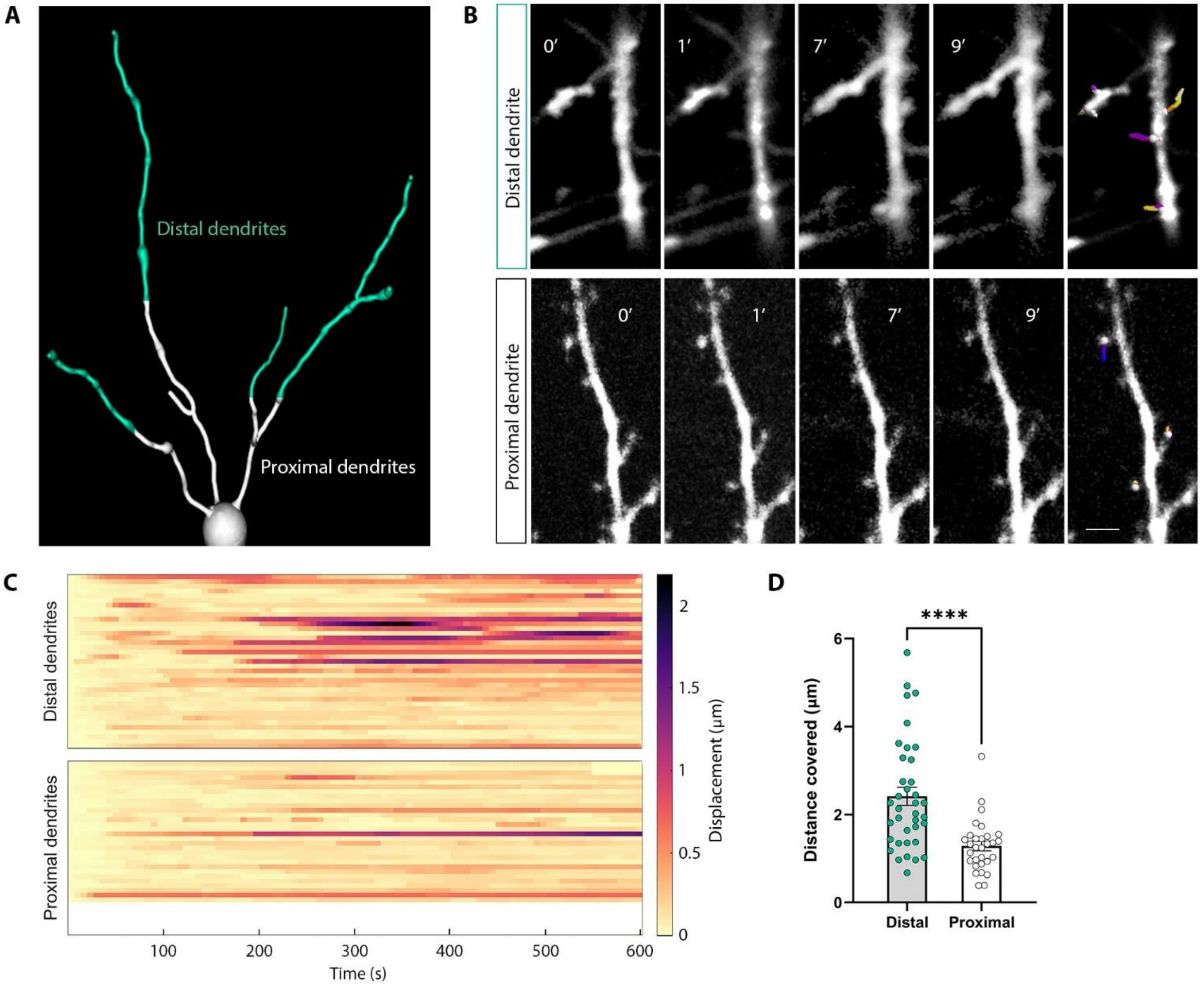


**Fig. S5. Dendrites display motile structural dynamics early in development.**
(**A**) Schematic of the basal arbor of a CA1 PN expressing tdTomato. Distal and proximal dendrites were continuously imaged for 10 minutes and are shown in cyan and white, respectively. (**B**) 2-photon time-lapse images of distal (top panel) and proximal dendrites (lower panel). Changes in dendritic arbor are indicated by colored tracks in far right images. Scale bar, 2 μm. (**C**) Raster plot of dendritic displacement traces (n = 37 distal dendrites, 10 cells from 4 mice; n = 30 proximal dendrites, 9 cells from 3 mice). (**D**) Comparison of the total distance covered of each distal and proximal dendrite that was tracked. ****$P < 0.0001$ (Mann-Whitney test).

**Fig. S6.**

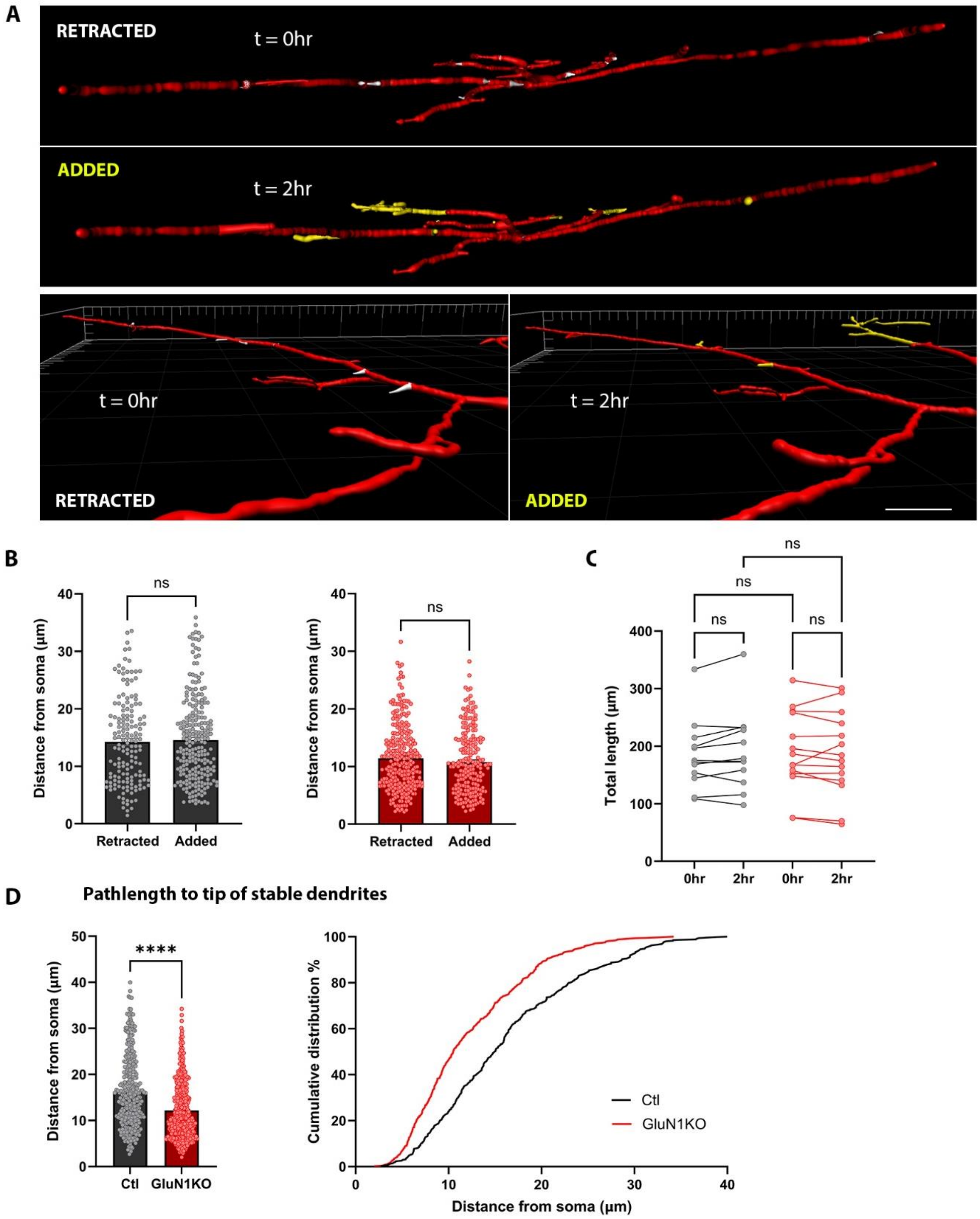


**Fig. S6. NMDARs contribute to dendritic complexity *in vivo*.**
(**A**) 3D reconstructions and side view visualizations of CA1 PN basal arbors showing retracted (white) and added (yellow) dendrites. Scale bar, 5 µm. (**B**) Comparison of pathlengths from the soma to the start point of retracted and added dendrites in Ctl neurons (n = 158 retracted, n = 241 added, 12 cells from 5 mice). $P = 0.7567$ (Mann-Whitney test). Comparison of pathlengths from

the soma to the start point of retracted and added dendrites in GluN1KO neurons (n = 270 retracted, n = 201 added, 14 cells from 5 mice). $P = 0.2392$ (Mann-Whitney test). (**C**) Total length measurements of Ctl (gray) and GluN1KO (red) basal arbors at timepoints 0 hr and 2 hr (n = 12 Ctl cells from 5 mice; n = 14 GluN1KO cells from 5 mice). $P > 0.9999$ each (Kruskal-Wallis test). (**D**) Measurement and cumulative distribution of pathlengths from the soma to the tip of stable dendrites of all neurons (n = 348 Ctl lengths, 12 cells from 5 mice; n = 495 GluN1KO lengths, 14 cells from 5 mice). ****$P < 0.0001$ (Mann-Whitney test).

**Fig. S7.**

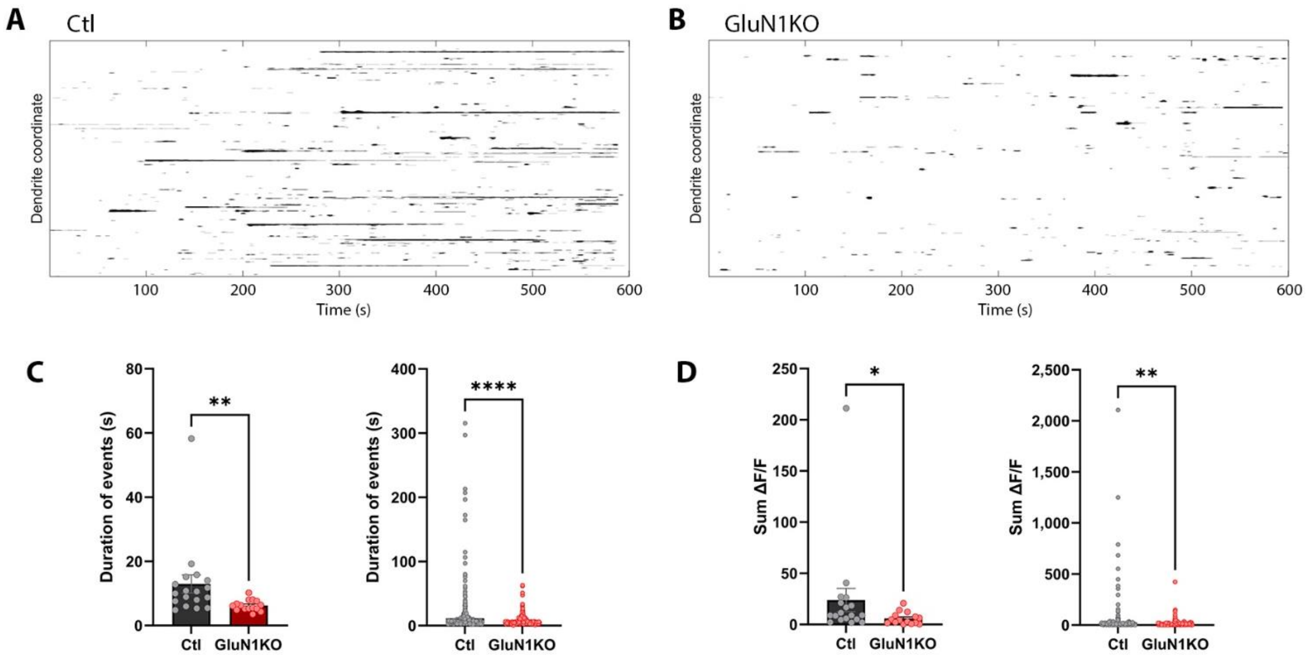


**Fig. S7. NMDARs give rise to long calcium transients during development.**
(**A, B**) Binary raster plots displaying events for Ctl cells and GluN1KO cells (1,826 Ctl dendrite coordinates, 18 cells from 3 mice; 1,765 GluN1KO dendrite coordinates, 15 cells from 2 mice). (**C**) Mean duration by cell and duration of all events (n = 779 Ctl events, 18 cells from 3 mice; n = 364 GluN1KO events, 15 cells from 2 mice). $**P = 0.0016$ and $****P < 0.0001$ (Mann-Whitney tests). (**D**) Mean sum ΔF/F by cell and sum ΔF/F of all events, normalized by dendritic length (n = 779 Ctl events, 18 cells from 3 mice; n = 364 GluN1KO events, 15 cells from 2 mice). $*P = 0.0147$ and $**P = 0.0038$ (Mann-Whitney tests).

Fig. S8.

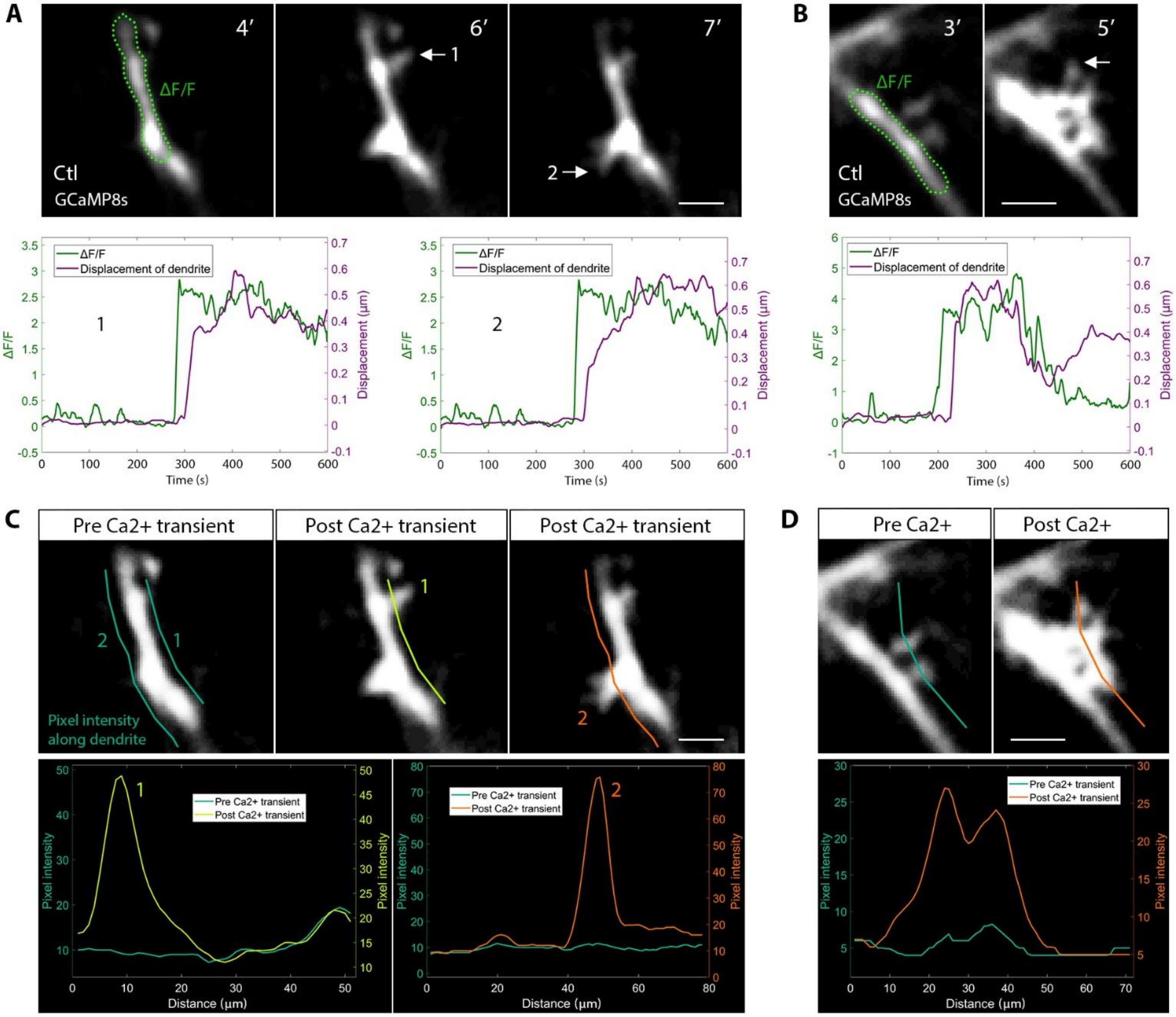


**Fig. S8. Additional Ctl cells showing structural changes coupled to long calcium transients.** (**A, B**) 2-photon time-lapse images of dendrites from two Ctl cells. Green demarcations outline ΔF/F ROIs, and white arrows indicate newly formed filopodia. ΔF/F an d dendritic displacement traces are overlaid to illustrate the temporal relationship between calcium transient and filopodial growth. Scale bar, 1 µm. (**C, D**) Line intensity scanning along dendrites to capture structural changes in dendritic arbor (see figure legend to Fig. 6B). Scale bar, 1 µm.

**Fig. S9.**

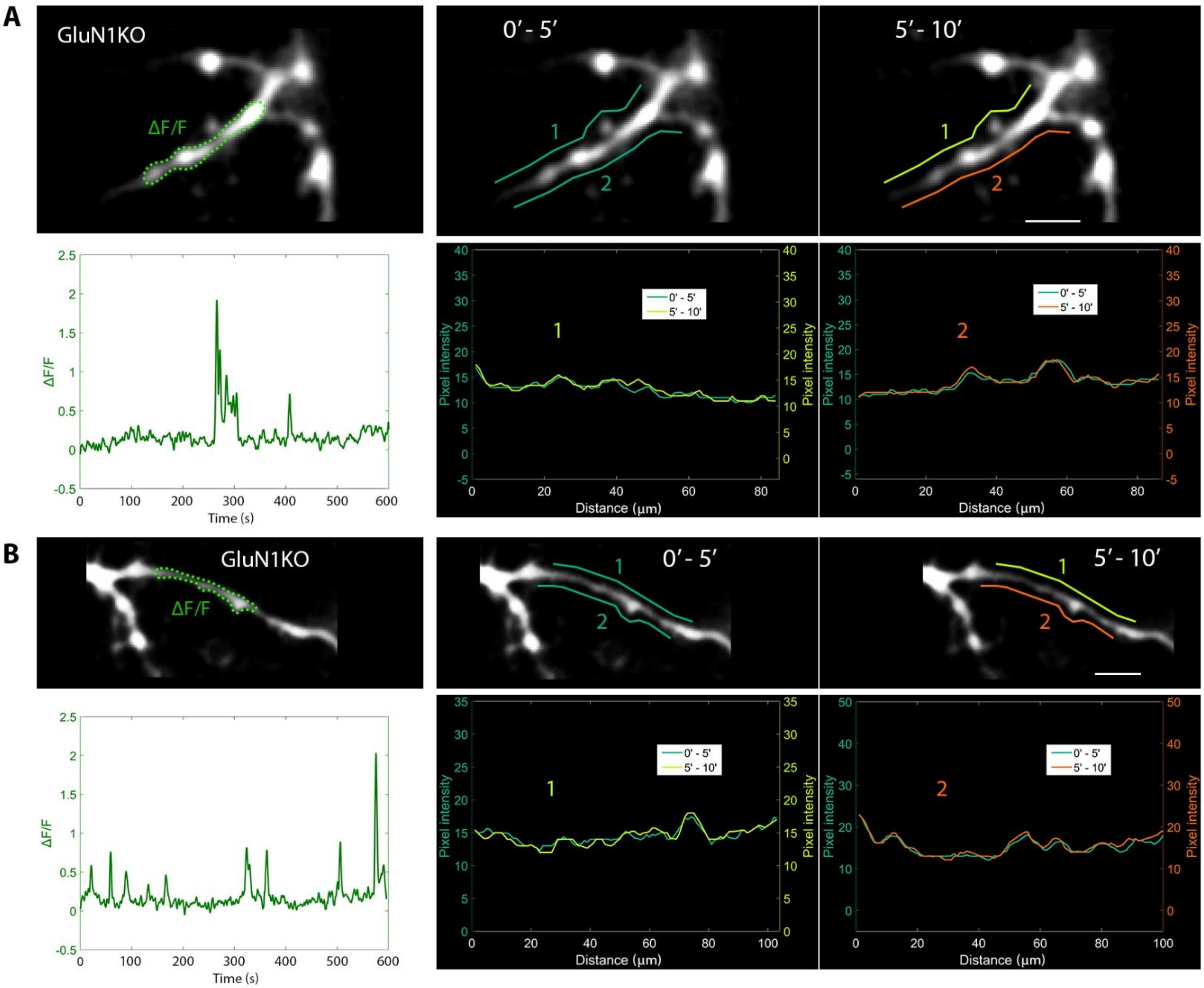


**Fig. S9. Lack of structural changes following calcium transients in GluN1KO cells.**
(**A, B**) Time-lapse images of dendrites from two GluN1KO cells. The ΔF/F ROI is outlined by the dashed green line. Pixel intensity measurements of lines drawn along dendrites in the first half (0’ – 5’) and second half (5’ – 10’) of imaging are overlaid to capture changes in dendritic arborization. Scale bar, 1 µm.

**Fig. S10.**

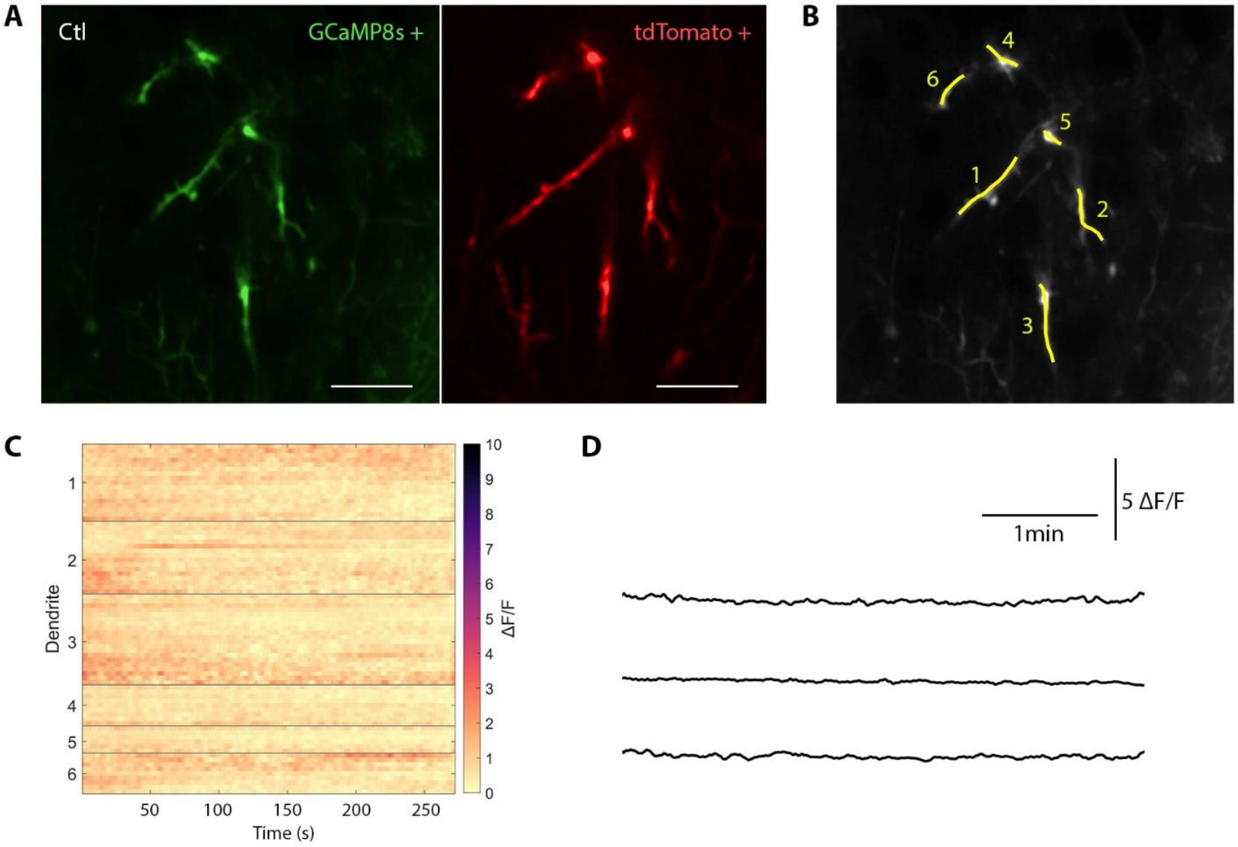


**Fig. S10. Absence of calcium transients when co-expressing GCaMP8s and tdTomato.**
(**A**) 2-photon images of an example Ctl CA1 PN expressing both GCaMP8s and tdTomato at P3. Scale bar, 5 µm. (**B**) 2D reconstructions of dendrites. (**C**) Raster plot depicting changes in fluorescence (ΔF/F) at coordinates along concatenated dendrites. (**D**) Exemplary ΔF/F traces from coordinates of 3 dendrites.

**Movie S1.**

**Movie S1.** Filopodial growth following a prolonged calcium transient 1. Frame-averaged movie depicting changes in dendritic structure correlating with a prolonged calcium transient. The cell is expressing GCaMP8s and is analyzed in Fig. 6A (1 s movie = 60 s real time).

**Movie S2.**

**Movie S2.** Filopodial growth following a prolonged calcium transient 2. Frame-averaged movie depicting changes in dendritic structure correlating with a prolonged calcium transient. The cell is expressing GCaMP8s and is analyzed in Fig. S7A (1 s movie = 60 s real time).

**Movie S3.**

**Movie S3.** Filopodial growth following a prolonged calcium transient 3. Frame-averaged movie depicting changes in dendritic structure correlating with a prolonged calcium transient. The cell is expressing GCaMP8s and is analyzed in Fig. S7B (1 s movie = 60 s real time).

**Movie S4.**

**Movie S4.** Lack of filopodial growth in a GluN1KO dendrite. Frame-averaged movie depicting an exemplary GluN1KO cell displaying dendritic calcium transients. No observable changes in dendritic structure are observed. The cell is expressing GCaMP8s and is from Fig. 6D (1 s movie = 60 s real time).
